# Increased mesoscale diffusivity in response to acute glucose starvation

**DOI:** 10.1101/2023.01.10.523352

**Authors:** Ying Xie, David Gresham, Liam Holt

**Affiliations:** New York University, School of Medicine, Institute for Systems Genetics, New York, USA; New York University, Center for Genomics and Systems Biology, Department of Biology, New York, USA

## Abstract

Macromolecular crowding is an important parameter that impacts multiple biological processes. Passive microrheology using single particle tracking is a powerful means of studying macromolecular crowding. Here we monitored the diffusivity of self-assembling fluorescent nanoparticles (μNS) in response to acute glucose starvation. mRNP diffusivity was reduced upon glucose starvation as previously reported. In contrast, we observed increased diffusivity of μNS particles. Our results suggest that, upon glucose starvation, mRNP granule diffusivity may be reduced due to changes in physical interactions, while global crowding in the cytoplasm may be reduced.

## Description

The cytoplasm is highly crowded, macromolecules such as RNA, protein and ribosomes occupy up to 30-40% of the cytoplasmic volume (Zhou et al., 2008; Zimmerman & Trach, 1991), leading to a phenomenon called macromolecular crowding. For simplicity, we will refer to macromolecular crowding as “crowding” henceforth. Crowding can influence biochemical reaction rates in several ways. First, high concentrations of crowders limit the excluded volume available to proteins or RNAs, thus impacting their thermal stability or proper folding (Pastore & Temussi, 2022; Smith et al., 2015).

Second, the crowded cytoplasm can favor intermolecular assembly, thereby accelerating reaction rates (Ellis, 2001; Rivas & Minton, 2018). Third, excessive crowding can dramatically decrease biomolecular diffusion rate, which slows down biochemical reactions (Ellis, 2001; Rivas & Minton, 2018). Thus, various cellular processes can be affected due to the change of biomolecular diffusion rate. For instance, actin polymerization is diffusion-limited and accelerated by increasing crowding (Drenckhahn & Pollard, 1986). Transcription activity partly depends on how fast a transcription factor diffuses to the gene loci and interacts with the enhancer(s) region (Berg et al., 1981). Therefore, changes in crowding can have profound effects on cell physiology.

The physical effect of crowding varies depending on the size of biomolecules. Length scales larger than individual macromolecular complexes such as ribosomes, but less than the size of the cell define the mesoscale (∼20 nm - 1 μm, Ekman et al., 2017). Many essential macromolecular complexes are of mesoscale size. DNA, RNA and proteins self-organize into a wide variety of structures such as mRNP granules and long range networks that are often highly dynamic (**Figure 1A**). Many mesoscale complexes are dynamically formed and remodeled depending on signals and environmental cues, and these processes typically involve active processes that consume energy (Sear et al., 2015). The biophysical principles behind tuning mesoscale complexes remain largely unknown. Modulating the crowding level may be a mechanism for cells to regulate mesoscale complexes. For example, ribosomes are the main crowders in the cytoplasm and their concentration strongly affects the diffusivity of 40 nm particles, and manipulating the ribosome concentrations through mTORC1 signaling has been shown to influence the phase separation behavior of synthetic condensates (Delarue et al., 2018).

**Figure 1.**
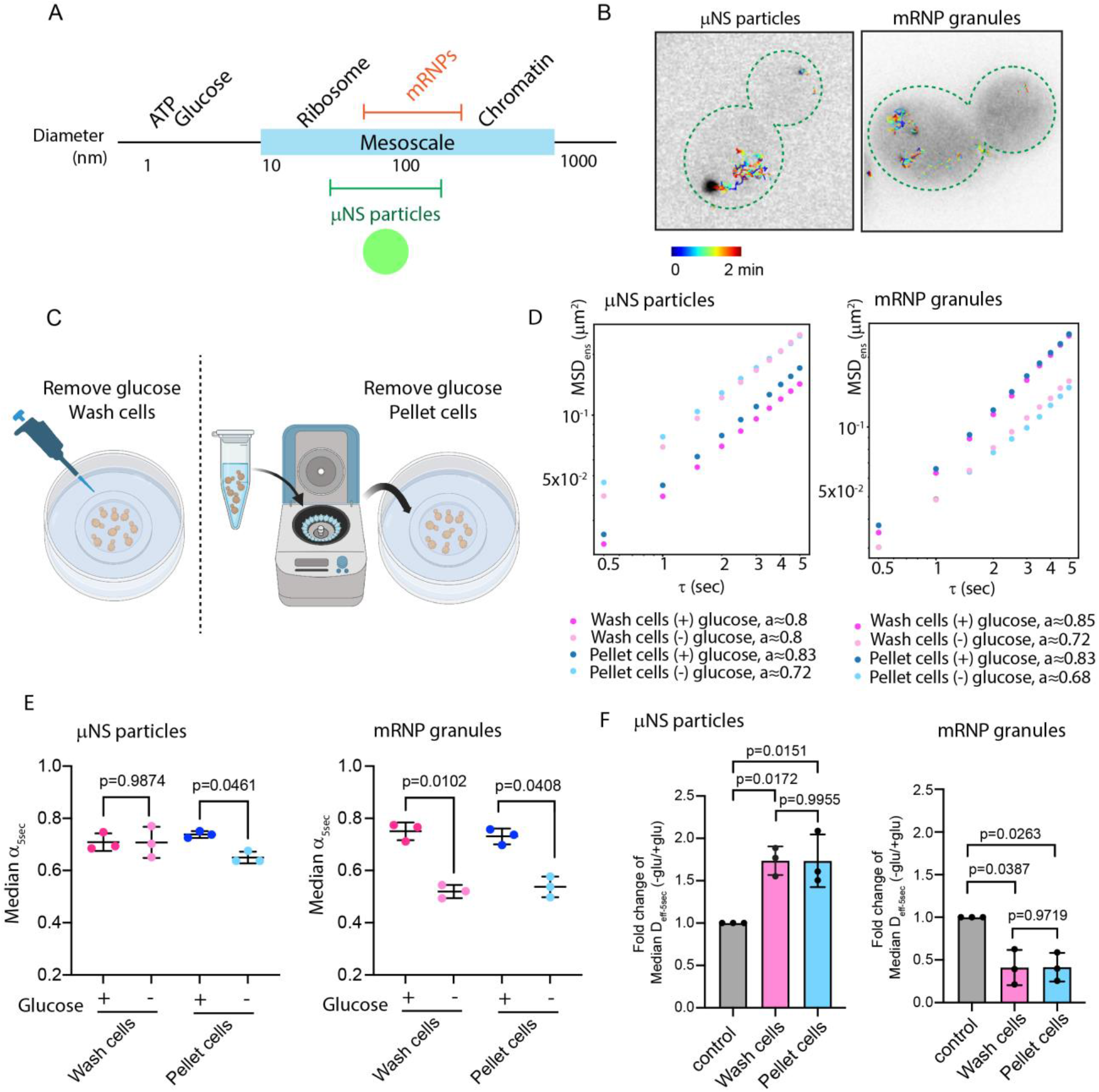
Mobility of μNS particles and mRNP granules upon acute glucose starvation. **A.)** The size of µNS and mRNP particles in relationship to other macromolecules. **B.)** Representative time projections of µNS particles and mRNP granules using live cell imaging in *Saccharomyces cerevisiae*. **C.)** Two methods used to perform acute glucose starvation. **D.)** Ensemble-averaged mean-squared displacement (MSD) versus time delay (ꚍ), log10 scale. A linear model was fitted to determine the anomalous exponent α values for each condition at 30 min.(n=700-1000 trajectories for µNS, n=300-500 trajectories for mRNP granules) **E.)** Median anomalous exponent α-5sec from individual trajectory of three biological replicates. (mean ± S.D., P values were determined from a paired two-tailed t test). **F)** Fold change of median effective diffusion coefficients at 5 sec (D_eff-5sec_) from 3 biological replicate experiments (mean ± S.D., P values were determined from a paired two-tailed t test).

Passive microrheology has been used in different studies to probe crowding levels and other cellular biophysical properties (Delarue et al., 2018; Etoc et al., 2018; Munder et al., 2016; Parry et al., 2014; Sabri et al., 2020; Shu et al., 2022). This method relies on analyzing the motions of tracer particles to infer the properties at the specific compartment. One such tracer particle, μNS, is derived from the mammalian orthorerovirus factory protein. The carboxyl-proximal regions (residues 471-721) of μNS fused with a GFP fluorescence tag can be easily expressed in cells through genetic engineering. μNS self-assembles into particles of varying size (50-150 nm) useful to probe mesoscale biophysical properties in the cell (Parry et al., 2014).

Prior work in bacteria showed that ATP depletion using toxins drastically reduces μNS particle diffusivity, suggesting that metabolic activity plays an important role in fluidizing the cytoplasm. Interestingly, this metabolic crisis showed a much stronger impact on the mobility of mesoscale particles > 50 nm in diameter, but had little effect on smaller particles (Parry et al., 2014). μNS particles have also been constructed in *Saccharomyces cerevisiae* (budding yeast) and *Schizosaccharomyces pombe* (Munder et al., 2016). Similar ATP depletion experiments in yeast cells balanced with different pH media suggests that acidification of the cytoplasm alters the biophysical properties of the cytosol. One possible explanation proposed by the study is that the acidified intracellular environment is close to the isoelectric point of a large portion of proteins, which leads to the formation of macromolecular assembly and therefore triggers the solid-like state transition of the cytoplasm. Interestingly, such solidification of cytoplasm correlates with a protective state for cell viability (Munder et al., 2016).

Chemically-induced ATP depletion is a relatively extreme scenario to challenge cell physiology. By contrast, microbes encounter frequent changes in nutritional status. For instance, glucose is an essential food source for budding yeast that can vary in abundance. Reorganization of intracellular structures and processes can be quickly and actively regulated upon nutrient perturbation coupled with the sudden change of physicochemical parameters, such as a drop in intracellular pH level, which was shown to actively signal the remodeling of the transcription program (Gutierrez et al., 2022; Triandafillou et al., 2020). Multiple processes including protein translation, endocytic trafficking, and inter organelle communications are rapidly re-organized upon glucose starvation in yeast (Janapala et al., 2019; Laidlaw et al., 2021; Wood et al., 2020). Such reorganization is likely to affect the crowding level. However, it remains unclear how crowding changes at the mesoscale when cells are challenged by external stresses.

A study analyzing the mobility of cytosolic mRNP granules upon acute glucose starvation has suggested a reduction of mesoscale particle mobility (Joyner et al., 2016). One possible explanation proposed by this study is that reduction of the intracellular volume coupled with an increase of crowding reduces mesoscale motion. However, changes in crowding might not be the only explanation for reduced mRNP granule dynamics. We hypothesize that increased physical interactions between mRNP granules and intracellular components might also constrain mobility. Therefore, we compared mRNP granules to μNS particles to understand how endogenous and exogenous probes move within the cytoplasm in response to acute glucose starvation (**Figure 1B**). Here, we study one of the representative mRNP granules previously used (Joyner et al., 2016), the *GFA1* transcript, an essential gene involved in chitin biosynthesis. In brief, the 24-PP7 stem-loops were integrated into the 3’ UTR of *GFA1* gene and the visualization of mRNPs was achieved by co-expressing the coat-binding protein, CP-PP7-3xYFP.

We also precisely controlled for several variables that are sometimes overlooked in acute glucose starvation experiments. First, crowding is sensitive to osmotic perturbation. The sudden removal of 2% glucose would reduce the osmotic pressure, leading to water influx and cell volume increase, thus complicating the interpretation of crowding changes. To avoid this, was osmotically balanced our starvation media with 2% sorbitol, which cannot be easily metabolized by budding yeast. Second, different groups have used different methods to switch cells to starvation medium. One method is to adhere cells on the imaging chamber and gently wash cells with starvation medium *in situ*. Another method is to use centrifugation to pellet then wash cells, typically 3-4 rounds of centrifugation/washing are used (Joyner et al., 2016). We were concerned that multiple centrifugation steps might contribute to additional compressive stress in cells (Peterson et al., 2012). As a result, we compared these methods to study both μNS particles and mRNP granules’ behavior upon acute glucose starvation (**Figure 1C)**.

## Results

To quantify the changes of diffusivities of μNS particles and mRNP granules, we plotted ensemble-averaged mean square displacements (MSDs) versus time delay (τ), with fits to determine anomalous exponent α values (**Figure 1D**). And we also calculated α values based on individual trajectories (**Figure 1E**). Both ensemble-time averaging analysis and individual trajectory analysis show consistent behavior for μNS particles and mRNP granules, displaying subdiffusive behavior (α < 1) (**Figure 1D and 1E**). The interpretation of the α value is very difficult but possible reasons include: local caging effect or non-specific interactions between the particles and subcellular structures. Assuming the μNS particles do not have regulated interactions, the subdiffusive behavior at proliferating cells is likely due to the local caging from crowders or subcellular structures. Notably, glucose starvation significantly reduced the α value (P value < 0.05) for mRNP granules in both starvation methods (**Figure 1E**), we reason that might be due to an increased physical interaction between mRNP granules and the cytosolic components.

Furthermore, we used hundreds of individual traces to quantify average displacement per unit time and computed effective diffusion coefficients at the 5 second timescale (D_eff-5sec_). A relative fold change of median D_eff-5sec_ in response to glucose starvation conditions to the median D_eff-5sec_ in proliferating cells was computed. Importantly, there was no obvious difference in our results for these two different methods used for inducing glucose starvation (**Figure 1F**). Consistent with the previous report (Joyner et al., 2016), a twofold decrease of diffusion rate was seen in mRNP granules (**Figure 1F**). By contrast, a 1.7 fold increase in diffusion rate was observed for μNS particles, consistent with macromolecular decrowding (**Figure 1F**).

## Discussion

A previous report has suggested caution in the interpretation of mRNP granule dynamic when using the PP7 stem–loops (PP7SL) system in budding yeast cells. Comparison with single-molecule fluorescence in situ hybridization (smFISH) revealed that engineering of the *GFA1* transcripts at 3’ UTR with stem-loops caused aberrant sequestration of the transcripts or the coat-binding proteins into processing bodies upon glucose starvation (Heinrich et al., 2017). Thus, it is likely that physical interactions of *GFA1-PP7SL* transcripts with RNA binding proteins in the processing bodies constrain diffusivity, leading to the observed reduction in D_eff-5sec_ for these mRNP granules upon glucose starvation. On the other hand, an increase of D_eff_ for μNS particles upon glucose starvation could be for several reasons: a reduction of crowder numbers or a restructuring of crowders may globally reduce the confinement of mesoscale macromolecules, or there could be an increase in non-thermal energy in the cell. Since ATP levels are decreased upon glucose starvation (Joyner et al., 2016), we do not favor this latter hypothesis. The contrasting changes of D_eff_ from μNS particles and mRNP granules highlight the importance of utilizing different foreign tracer probes to study the biophysical properties of the cell.

## Materials and Methods

### Yeast culture

*Saccharomyces cerevisiae* strains used in this study are listed in Table 1. Strains were revived from -80°C freezer on YPD plate for an overnight growth, and then a patch of cells were inoculated in synthetic complete media + 2% dextrose (SCD) according to standard Cold Spring Harbor Protocols (“Synthetic Complete (SC) Medium,” 2016). The cultures were grown at 30°C in a rotating incubator for 4-5 hours without exceeding an O.D.600 of 0.4. Afterwards, the cultures were diluted for an overnight growth in order to reach O.D.600 between 0.1 and 0.4 for the next day ‘s imaging experiment.

**Table 1.**
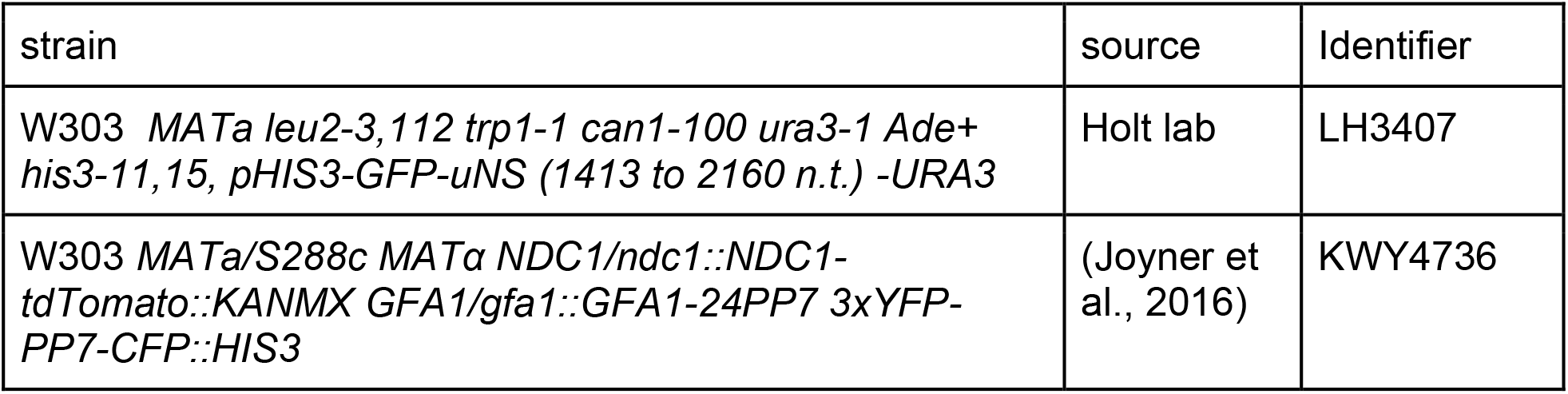
Yeast strains used in this study

### Acute glucose starvation

Switching cells to starvation medium without pelleting by centrifugation: Cells with culture medium reaching O.D.600 = 0.1–0.4 were applied to imaging chamber, which was pre-coated by 1mg/ml concanavalin A to immobilize cells on the coverslip. After waiting for 5-10 min for cells to settle down on the coverslip, the culture medium was removed completely, and four additional wash of cells with SCD or SC medium supplemented with 2% sorbitol (to balance the osmotic pressure) was performed. Cells were imaged at 30 min afterwards.

Switching cells to starvation medium by centrifugation: 1 ml cells grown in SCD media (O.D.600 = 0.1–0.4) were collected by centrifugation (3000 rpm) for 2 min. The supernatant was removed and cells were resuspended in 1 ml of SC with 100 mM sorbitol media. Four additional wash steps followed, with two 2 min spins (6000 rpm) succeeded by two 1 min spins. The cells were suspended into fresh SC+2% sorbitol or SCD media for analysis at 30 min.

### Imaging and particle tracking

Single particle tracking in *Saccharomyces cerevisiae* was performed for the uNS particles and GFA1-mRNP granules. The particles were imaged using Andor Yokogawa CSU-X confocal spinning disc on a Nikon Ti2 Eclipse microscope and fluorescence was recorded with a sCMOS Prime 95B camera (Photometrics) with a 63x objective (pixel size: 0.18 μm), at a 500 ms image capture rate, with a time step for 1 min. The tracking of particles was performed with the Mosaic suite of FIJI (Sbalzarini & Koumoutsakos, 2005), using the following typical parameters: radius = 2; cutoff = 0; percent: variable, a link range of 1, and a maximum displacement of 5 pixels, assuming Brownian dynamics.

### Quantification of mesoscale rheology

For every trajectory, the time-averaged mean-square displacement (MSD) was quantified as defined in (Delarue et al., 2018; Joyner et al., 2016; Munder et al., 2016; Shu et al., n.d.2022). To characterize individual particle trajectories, we calculated effective diffusion coefficients by fitting MSD with a linear (diffusive) time dependence within ten time points. Particle trajectories with more than ten time points were selected for analysis to reduce tracking error. Time-averaged MSD for each trajectory is fitted using a linear time dependence within first ten time intervals:

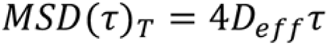

where D_eff_ is the effective diffusion coefficient for each trajectory, τ is the time delay. And we refer to the median value D_eff_ from all the trajectories at each condition to represent its effective diffusion coefficient.

Ensemble-time averaged MSD was applied for better indication of anomalous exponent α at each condition. which is ensemble-averaged from all time-averaged MSD for trajectories with more than ten time points:

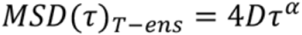

All quantifications were performed in the programme GEMspa (https://github.com/liamholtlab/GEMspa) and graph plots were generated in GraphPad Prism 9.

## Acknowledgements

Support from NIH to DG (R01-GM134066, R01-GM107466) L.J.H. was funded by NIH R01 GM132447 and R37 CA240765, the American Cancer Society, the NIH Director’s Transformative Research Award TR01 NS127186, the Air Force Office of Scientific Research (AFoSR FA9550-21-1-3503 0091), and the Human Frontier Science Program (RGP0016/2022-102)

